# An optimized ribodepletion approach for *C. elegans* RNA-sequencing libraries

**DOI:** 10.1101/2021.01.04.425342

**Authors:** Alec Barrett, Rebecca McWhirter, Seth R Taylor, Alexis Weinreb, David M Miller, Marc Hammarlund

**Affiliations:** Department of Genetics, Yale University School of Medicine, New Haven, CT; Department of Cell and Developmental Biology, Vanderbilt University School of Medicine, Nashville, TN; Program in Neuroscience, Vanderbilt University School of Medicine, Nashville, TN, USA; Department of Neuroscience, Yale University School of Medicine, New Haven, CT

## Abstract

A recent and powerful technique is to obtain transcriptomes from rare cell populations, such as single neurons in *C. elegans,* by enriching dissociated cells using fluorescent sorting. However, these cell samples often have low yields of RNA that present challenges in library preparation. This can lead to PCR duplicates, noisy gene expression for lowly expressed genes, and other issues that limit endpoint analysis. Further, some common resources, such as sequence specific kits for removing ribosomal RNA, are not optimized for non-mammalian samples. To optimize library construction for such challenging samples, we compared two approaches for building RNAseq libraries from less than 10 nanograms of *C. elegans* RNA: SMARTSeq V4 (Takara), a widely used kit for selecting poly-adenylated transcripts; and SoLo Ovation (Tecan Genomics), a newly developed ribodepletion-based approach. For ribodepletion, we used a custom kit of 200 probes designed to match *C. elegans* rRNA gene sequences. We found that SoLo Ovation, in combination with our custom *C. elegans* probe set for rRNA depletion, detects an expanded set of noncoding RNAs, shows reduced noise in lowly expressed genes, and more accurately counts expression of long genes. The approach described here should be broadly useful for similar efforts to analyze transcriptomics when RNA is limiting.

## INTRODUCTION

RNA sequencing (RNAseq) is a well-established method for assessing gene expression. In *C. elegans*, RNAseq can be combined with cell enrichment by fluorescence activated cell sorting (FACS) for studies of gene expression in individual cell types, enabling a high resolution view of cell type-specific transcription (1–5). Sequencing library construction is a significant variable in RNAseq experiments from FACS-enriched *C. elegans* samples, as well as from other samples with low amounts of input RNA. Optimizing library construction to maximize data recovery is therefore an important goal.

At approximately 90% of total RNA in most cells, the prevalence of ribosomal RNA (rRNA) constitutes a major challenge for RNAseq profiling of other RNA species (6). To efficiently sequence the remaining 10% of cellular RNA, rRNA is typically excluded during library construction. One common approach to meet this goal is the use of poly-d(T) primers to favor cDNA synthesis from poly-adenylated (polyA) RNA versus rRNA, which is typically not poly-adenylated. In an alternative strategy, known as ribodepletion, oligonucleotides complementary to specific rRNA sequences are used to deplete rRNA transcripts from the library by either bead affinity extraction (7, 8) or directed enzymatic cleavage (9). Overall, polyA approaches can be more efficient at excluding rRNA compared to ribodepletion strategies (10). However, polyA methods require the RNA input to be largely free from degradation, tend to bias coverage towards the 3’ end of transcripts and exclude ncRNA (noncoding RNA) species that lack polyA tails. In contrast, ribodepletion is better suited for low quality samples, as random primer amplification is more likely to capture fragmented RNAs. Importantly, ribodepletion preserves ncRNAs and allows library construction methods that favor more uniform gene body coverage (10–12).

Although recent advancements in library preparation methods have demonstrated that both polyA and ribodepletion methods are feasible for ultra-low input samples from *C. elegans* (1, 13), most published *C. elegans* studies have used polyA approaches (14–18), and a ribodepletion approach was found to retain high levels of rRNA in the final library (1), potentially because these rRNA ribodepletion oligonucleotides were optimized for mammalian rRNA.

Here, we introduce a ribodepletion approach that uses oligonucleotides specifically designed to match *C. elegans* rRNA sequences and test the idea that this optimized ribodepletion strategy can produce favorable results on low-input RNA samples. We compared polyA (SMARTseq V4) and ribodepletion (SoLo Ovation) approaches, using as input <10ng of total RNA prepared from FACS-enriched *C. elegans* neurons. We constructed multiple libraries with each strategy and sequenced the resulting libraries at high depth. Detailed comparison of the results indicated that although high-quality libraries can be obtained with either method, SoLo ribodepletion with *C. elegans* specific probes has significant advantages over the commonly used SMARTseq polyA approach, including increased detection of noncoding RNAs, reduced noise for lowly expressed genes, and more accurate counts for long genes.

## RESULTS

### Designing a custom set of *C. elegans* rRNA depletion probes

Ribodepletion strategies are sequence dependent, and mismatches or gaps between the oligonucleotide probe set and rRNA sequences may allow substantial rRNA sequences to remain in the finished sequencing library. rRNA depletion probe sets designed for mammalian samples perform poorly in *C. elegans* (1). We therefore designed a probe set to match *C. elegans* rRNA. We collected fasta files from all *C. elegans* rRNA genes in the WS235 genome assembly and removed duplicate sequences to identify 200 unique rRNA probes for use in our ribodepletion experiments (Methods).

### Collecting pan-neuronal samples via FACS

We tested different methods to optimize library construction from low amounts of RNA typically provided by FACS-sorted samples from *C. elegans*. We dissociated L4-stage larvae with SDS and protease treatments (1, 3, 4), and then used FACS to enrich for cells expressing the neuron-specific reporter *Prab-3*::RFP (19). For each RNA sample, ~25,000 cells were collected directly into TRIzol and RNA extracted by Direct-zol RNA purification (Zymo Research) columns. For this study, we used 4 independently generated RNA samples, with one of those samples split into two technical replicates during purification, one DNase pre-treated, and one not (see Methods, Supplementary table 1). Yields of total RNA ranged from 2.2 ng to 7.2 ng for each sample (Supplementary Table 1).

### SMARTseq excludes more rRNA whereas SoLo shows fewer duplicate reads

Next, for direct comparisons of polyA and ribodepletion approaches, we split each RNA sample in half, resulting in two sets of matched samples, with five total replicates in each. We built sequencing libraries for all samples, using either a polyA approach (SMARTseq V4) or a ribodepletion strategy (SoLo Ovation) for each pair of matched samples. Briefly, for SMARTseq libraries, cDNA was prepared with poly-d(T) primers and then amplified. Fragmentation, adapter ligation, and final library preparation were performed using the Illumina Nextera XT kit. For the SoLo approach, cDNA libraries were prepared using random primers prior to fragmentation, adapter ligation, and amplification. After isothermal amplification, the *C. elegans* custom probe set was used to direct cleavage of rRNA fragment adapters prior to a final round of amplification. We sequenced all libraries on an Illumina Hiseq 2500 machine with paired end 75bp reads to a depth of 15 to 37 million read pairs per library. After sequencing, fastq files were checked for read quality using fastQC. One SMARTseq sample failed quality control (low per base quality, and highly 3’ biased gene coverage), and this SMARTseq library and the corresponding SoLo library were removed from further analysis. Of the four paired sets of samples, SoLo sequenced libraries had an average of 17.96 million read pairs, and SMARTseq libraries had an average of 31.77 million read pairs (Supplementary Table 1).

We assessed the basic properties of the ribodepletion and polyA libraries using metrics that reflect the relative number of useful reads. We defined useful reads as those that are not PCR duplicates and that do not map to rRNA genes. We used STAR to map all reads to the WS235 genome with default parameters (20), and observed that SMARTseq and SoLo samples show similar mapping rates, averaging 67% and 59% respectively (Figure 1A). After mapping we marked and removed duplicate reads based on position using SAMtools (21). The percentage of reads remaining after deduplication provides a measure of duplicate reads. Using this metric, SoLo libraries had consistently lower rates of PCR duplicates (mean 33.7% reads remaining after deduplication) than SMARTseq libraries (mean 26.5% reads remaining after deduplication) (Figure 1B).

**Figure 1:**
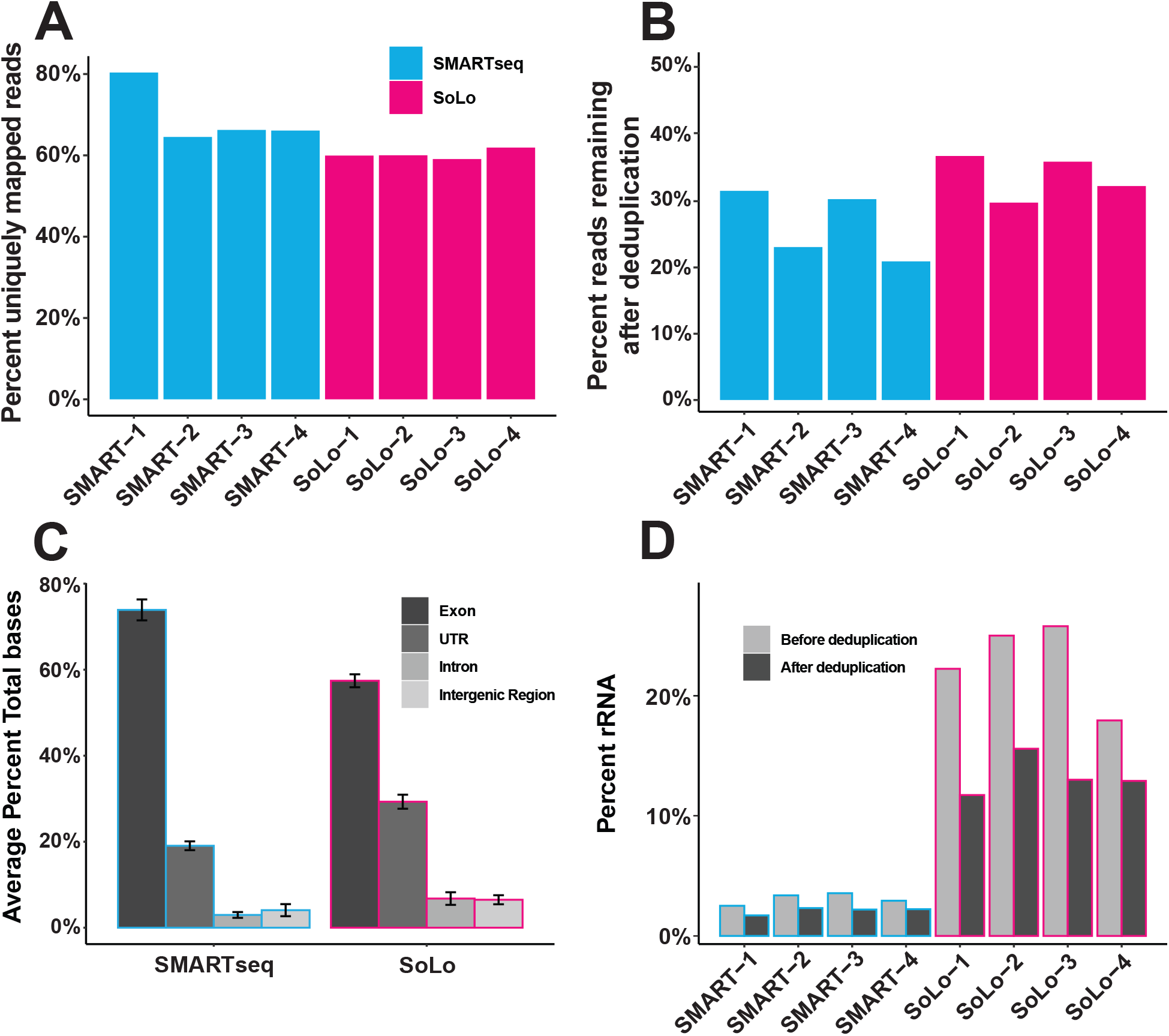
Mapping summary. A) Bar graph showing the percent of all reads mapped from raw fastq files for SMARTseq replicates (blue) and SoLo replicates (magenta). B) Bar graph showing the percent of mapped reads that were not marked as likely PCR duplicates. C) Bar graph showing the percent of bases that mapped to exons, UTRs, introns, and intergenic regions. Error bars show 95% confidence intervals. D) Bar graph showing the percent of mapped reads counted in rRNA genes before and after deduplication.

To assess where reads mapped, we compared the percentage of bases that mapped to exons, untranslated regions (UTRs), introns, and intergenic regions. SMARTseq libraries mapped an average of 73.9% of bases to exons, 19.0% to UTRs, 3.0% to introns, and 4.1% to intergenic regions. SoLo libraries mapped an average of 57.4% of bases to exons, 29.3% to UTRs, 6.8% to introns, and 6.5% to intergenic regions (Figure 1C). As UTRs are included in the gene models used for assigning counts, the average total percent of gene-feature mapped reads is 92.9% for SMARTseq and 86.7% for SoLo. Next, we assessed the fraction of rRNA reads in both original and deduplicated libraries. Prior to deduplication, SMARTseq samples had an average of 3.0% (range = 2.5% - 3.6%), while SoLo samples had an average of 22.7% rRNA reads (range = 17.9% - 25.7%). In deduplicated libraries, SMARTseq samples had an average of 2.1% (range = 1.7% - 2.3%), while SoLo samples had an average of 13.3% rRNA reads (range = 11.7% - 15.6%) (Figure 1D). Assuming rRNA is 90% of the cellular RNA, these data indicate that SMARTseq removed an average of 99.8% of the rRNA, whereas SoLo removed an average of 98.3% of the rRNA. Overall, these data indicate that both techniques result in efficient selection against rRNA but that the SMARTseq polyA-based approach performed better than SoLo in rRNA removal (Figure 1D).

### SoLo and SMARTseq detect largely overlapping gene sets

Next, we compared the overall number of expressed genes between ribodepletion and polyA libraries. Each sample was normalized using the GeTMM method, first to gene length and then to the Trimmed Mean of M-values (TMM) corrected library size to account for intra-sample and inter-sample variation (22). Average GeTMM values for all genes were generally correlated between SMARTseq and SoLo samples, with a Spearman correlation coefficient of 0.79. Within each technique, replicates were also highly correlated (Supplementary Figure 1A-C). We then calculated 95% confidence intervals for all genes within SMARTseq and SoLo samples. We defined expressed genes as those genes where the lower bound of the 95% confidence interval is > 5 GeTMM. Using this definition, we called 6,146 genes expressed in SMARTseq, and 7,108 genes expressed in SoLo. The majority of expressed genes (5,104) were called expressed in both approaches (Figure 2A). Similarly, we defined not-expressed genes as those genes where the upper bound of the confidence interval is < 5 GeTMM. The remaining genes, with confidence intervals that include 5 GeTMM, we consider to be genes for which expression is ambiguous. Interestingly, “ambiguous” genes were more common in SMARTseq samples (Figure 2B). Overall, ribodepletion and polyA priming approaches resulted in broadly similar results for gene expression. However, substantial differences exist between the two approaches, with 1,042 out of 6,146 expressed genes in SMARTseq not identified in SoLo, and 2,004 out of 7,108 expressed genes in SoLo not identified in SMARTseq (Fig 2A). The existence of these differences raises the question of how differences in RNA capture or amplification affect specific RNA types.

**Figure 2:**
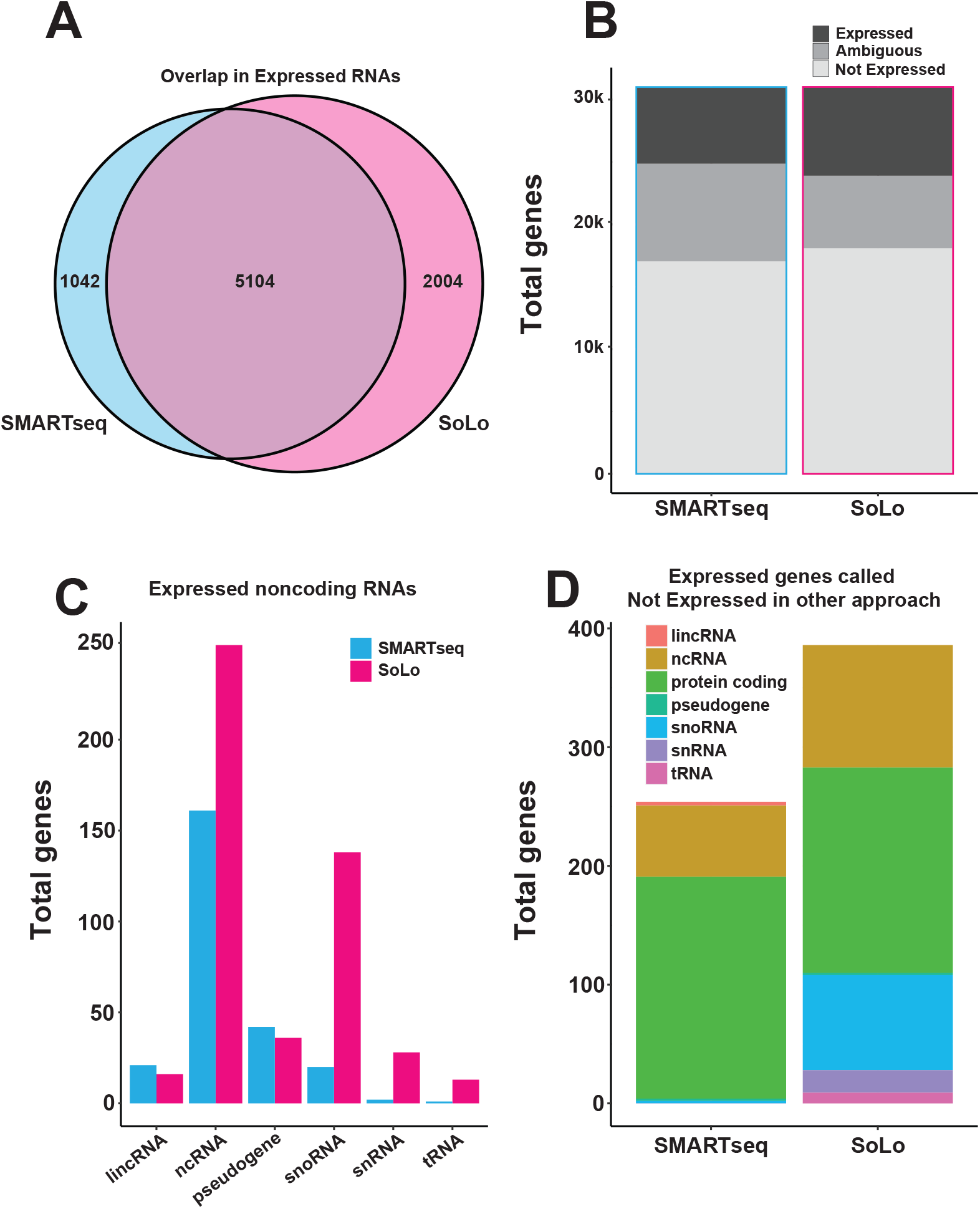
SoLo detects classes of noncoding RNAs missed in SMARTseq. A) Venn diagram showing the overlap between high confidence expressed genes in SMARTseq and SoLo. B) Bar graph showing gene detection for SMARTseq and SoLo using all genes, with three levels based on confidence intervals (CI): expressed (lower bound of CI > 5 GeTMM), “ambiguous” (CI overlaps 5 GeTMM), and “not expressed” (upper bound of CI < 5 GeTMM). C) Bar graph showing the number of noncoding RNAs called expressed (lower bound of CI > 5 GeTMM) in SMARTSeq (blue) and SoLo (magenta), in separate categories for each RNA type. D) Bar graph showing genes called “expressed” (lower bound of CI > 5 GeTMM) in one technique, and “not expressed” (upper bound of CI < 5 GeTMM) in the other, broken down by gene type.

### SoLo detects an expanded set of noncoding RNA species

A potential source of differences between the datasets is the role of polyA tails in cDNA synthesis. Many noncoding RNAs lack 3’ polyA tails and are thus unlikely to be efficiently captured by SMARTseq cDNA synthesis, which depends on poly-d(T) priming. To test for this possibility, we compared the detection rates of six classes of noncoding RNAs between SoLo and SMARTseq, using the 5 GeTMM threshold for calling genes expressed. Of these classes, pseudogenes and long intergenic noncoding RNAs (lincRNAs) often have polyA tails (23, 24), and we found that these classes are detected at similar frequencies between SoLo and SMARTseq (Figure 2C). By contrast, small nuclear RNAs (snRNAs), small nucleolar RNAs (snoRNAs), and transfer RNAs (tRNAs) usually lack polyA tails (25), and we found that these classes are detected at much higher frequencies in SoLo than SMARTseq libraries (Figure 2C). In comparison to SMARTseq, SoLo calls 6.9 times as many snRNAs “expressed”, 24 times as many snoRNAs, and 13 times as many tRNAs (Figure 2C). Additionally, of the genes called “expressed” by SoLo, 57.6% of snRNAs, 33.3% of snoRNAs, and 82.2% of tRNAs have zero counts in any SMARTseq replicate. A final class of RNAs, uncategorized noncoding RNAs (ncRNAs), was detected at high levels in both approaches, although SoLo detects ~50% more ncRNA transcripts, suggesting that this category contains a mix of poly-adenylated and non-poly-adenylated transcripts.

To determine the contribution of noncoding RNAs to discrepancies in gene detection between techniques, we focused in on the genes that are confidently called “expressed” in one technique, but “not expressed” in the other, not considering “ambiguous” genes. Breaking down those sets by biotype, we see that 67 (31.5%) of the 254 SMARTseq “expressed” exclusive genes are noncoding RNAs, whereas 213 (55.1%) of the 386 SoLo “expressed” exclusive genes are noncoding RNAs (Figure 2D). This analysis indicates that a majority of the genes confidently detected by the SoLo method but not by SMARTseq are noncoding RNAs.

### SoLo samples show reduced variance among lowly expressed protein coding genes

SMARTseq has many more genes for which expression was “ambiguous” than SoLo (Figure 2B). These data could indicate a difference in noise between the techniques, which might account for some of the remaining differences in apparent gene expression that are not explained by differences in noncoding RNA detection. To explore this possibility, we generated GeTMM values using only counts that map to protein coding genes. Using the thresholding method described above, SMARTseq libraries yielded 5,899 “expressed” genes, 8,320 “not expressed” genes, and 6,288 “ambiguous” genes. SoLo libraries yielded 6,625 “expressed” genes, 10,091 “not expressed” genes, and 3,731 “ambiguous” genes (Figure 3A). Thus, SoLo libraries yielded more protein coding genes that are confidently called either “expressed” or “not expressed”, whereas SMARTseq showed almost twice as many “ambiguous” genes.

**Figure 3:**
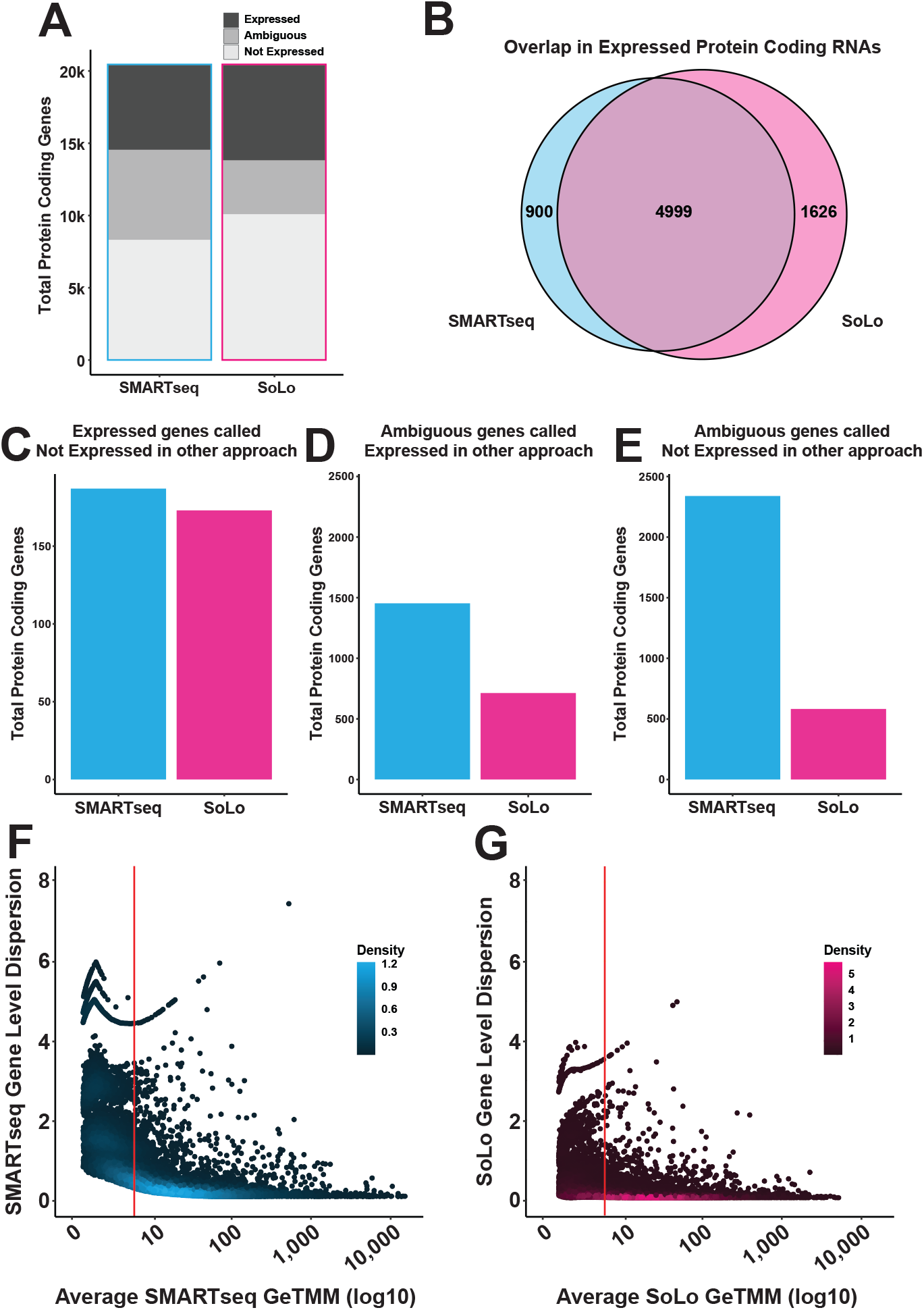
SMARTseq protein coding genes show higher dispersion. A) Venn diagram showing the overlap between “expressed” protein coding genes in SMARTseq and SoLo. B) Bar graph showing gene detection for SMARTseq and SoLo using protein coding genes, with three levels based on confidence intervals (CI): “expressed” (lower bound of CI > 5 GeTMM), “ambiguous” (CI overlaps 5 GeTMM), and “not expressed” (upper bound of CI < 5 GeTMM). C) Bar graph showing the number of protein coding genes called “expressed” (lower bound of CI > 5 GeTMM) in one technique and “not expressed” (upper bound of CI < 5 GeTMM) in the other. D) Bar graph showing the number of protein coding genes called “ambiguous” (CI overlaps 5 GeTMM) in one technique and “expressed” (lower bound of CI > 5 GeTMM) in the other. E) Bar graph showing the number of protein coding genes called “ambiguous” (CI overlaps 5 GeTMM) in one technique and “not expressed” (upper bound of CI < 5 GeTMM) in the other. F) Scatter plot showing the edgeR gene level dispersion estimate against average log10 GeTMM levels for SMARTseq protein coding genes (average SMARTseq GeTMM > 0.5). Red line shows 5 GeTMM. G) Scatter plot showing the edgeR gene level dispersion estimate against average log10 GeTMM levels for SMARTseq protein coding genes (average SoLo GeTMM > 0.5). Red line shows 5 GeTMM.

Other than this important difference, the results from the two techniques for protein coding genes were similar. Similar to results analyzing expression data for all genes (Figure 2B), the majority of protein coding genes that were confidently called “expressed” in either technique were called “expressed” in both techniques (Figure 3B). Additionally, mRNAs called “expressed” in one technique were very rarely called “not expressed” in the other (SMARTseq: 187, SoLo: 173) (Figure 3C). GeTMM values for protein coding genes were highly correlated between the two techniques, with a Spearman correlation coefficient of 0.88 (Supplementary Figure 1D). Among the 6,288 “ambiguous” SMARTseq genes, 1,453 (23.1%) were called “expressed” in SoLo, and 2,339 (37.2%) were called “not expressed” in SoLo. For SoLo samples, of the 3,731 “ambiguous” genes, 713 (19.1%) were called “expressed” in SMARTseq and 582 (15.6%) were called “not expressed” in SMARTseq. Overall numbers of “ambiguous” genes were higher in SMARTseq. (Figure 3D-E).

We considered the possibility that ambiguity in gene expression might correlate with lower gene expression, since lowly expressed genes might be more prone to noise. Dispersion is a measure of variance calculated when fitting expression data to a negative binomial model. We estimated the library-to-library dispersion of each gene within each technique using edgeR, and plotted these values against average gene expression. We observed that SMARTseq had more dispersion than SoLo across all GeTMM values. However, this difference is strongest among lowly expressed genes (Figure 3F-G, Supplementary Figure 2A). Comparing confidence interval size (another indicator of variance), on a gene-by-gene level, reveals a similar trend (Supplementary Figure 2B). These data suggest that SoLo produces consistent values for protein coding genes across a wider range of expression levels than SMARTseq, and that at least some of the difference in genes confidently called “expressed” or “not expressed” is explained by intra-technique variance among low expressed genes.

### SoLo shows enhanced detection of long genes

Besides noise, a potential source of the differences in protein coding gene expression between the techniques is bias that depends on gene length. In general, RNAseq expression analysis counts the number of reads per gene, which is dependent on the number of cDNA fragments from that gene in the sequencing library. Longer genes have the potential to be represented in the library by more fragments, and thus accumulate more reads than short genes with the same number of RNA molecules in the sample. Thus, read counts must be normalized to the known length of the transcript in the genome assembly—in essence, reads per gene are divided by transcript length. However, this normalization approach assumes that read abundance increases linearly with read length, across all lengths. Length-dependent bias can occur if, for example, reads are depleted from the 5’ end of long genes, but not short genes. In polyA-primed approaches such as SMARTseq, this depletion can occur due to RNA degradation, particularly of longer transcripts, or due to incomplete processivity during reverse transcription. To test the idea that gene length correlates with differences between the two approaches, we examined the gene length distribution for protein coding genes called “expressed” only in SMARTseq or SoLo and compared these unique genes to the distribution of gene lengths for all protein coding genes (Figure 4A). We found that SMARTseq exclusive genes (median 1.07kb) are generally very close to the distribution for all protein coding genes protein coding genes (median 1.09kb), whereas SoLo exclusive genes are enriched for longer genes (median 1.74kb) (SMARTseq p = 0.022; SoLo: p = 1.07 * 10^−15^). This analysis suggests that differences in library construction may result in length-dependent biases in the data.

**Figure 4:**
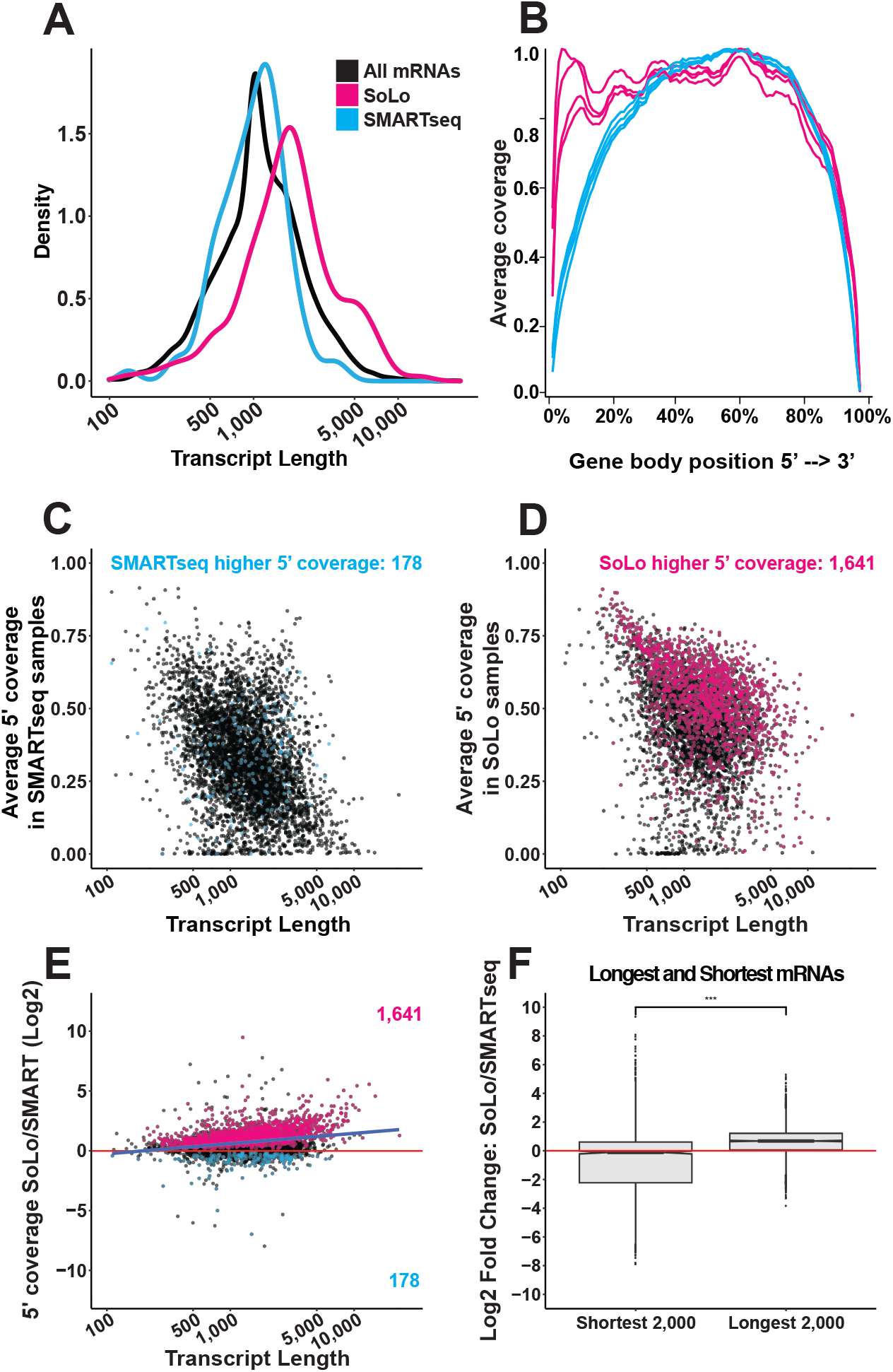
SoLo shows higher expression for long genes. A) Density graph showing the transcript length distribution for all protein coding mRNAs (black), mRNAs “expressed” (lower bound of CI > 5 GeTMM) in SMARTseq but “not expressed” (upper bound of CI < 5 GeTMM) in SoLo (blue), and mRNAs “expressed” in SoLo but “not expressed” in SMARTseq (magenta). B) Line plot showing the average normalized gene body coverage for all protein coding genes > 100bp. Left to right, 5’ to 3’. SMARTseq replicates shown in blue, Solo replicates shown in magenta. C) Scatterplot showing the average SMARTseq coverage of the 5’ end of all protein coding genes called “expressed” in both SMARTseq and SoLo. 178 genes found with significantly higher 5’ coverage in SMARTseq colored blue, paired t-test, BH adjusted p-value < 0.05. D) Scatterplot showing the average SoLo coverage of the 5’ end of all protein coding genes called expressed in both SMARTseq and SoLo. 1,641 genes found with significantly higher 5’ coverage in SoLo colored magenta, paired t-test, BH adjusted p-value < 0.05. E) Scatter plot showing the log2 fold ratio of SoLo and SMARTseq 5’ gene coverage for protein coding genes called expressed in both SMARTseq and SoLo. Significant genes called higher in SoLo colored magenta, genes called higher in SMARTSeq colored blue. Paired t-test, BH adjusted p-value < 0.05. Linear model: 0.872* log10(Length) - 2.084, R^2^ = 0.073. F) Box plot showing the edgeR log fold change for gene expression among the 2000 shortest protein coding genes (length > 100bp) and 2000 longest protein coding genes, SoLo/SMARTseq. Wilcoxon test, ***: p-value < 0.001.

As an additional test of this result, we examined read coverage across the length of each gene in all samples. Using the RSeQC suite to measure average coverage (26), we found that SMARTseq coverage drops off near the 5’ end. SoLo samples show no such drop-off and generally cover the entire length of the gene (Figure 4B). Given this difference, we hypothesized that longer genes may be especially prone to low 5’ end coverage in SMARTseq libraries. We defined the 5’ end as the first 20% of each gene and found that 5’ end coverage tends to decrease for both SMARTseq and SoLo as gene length increases, but that the drop-off is much steeper for SMARTseq (Figure 4C-D). Comparing the 5’ coverage at the gene level shows that 178 genes have higher coverage in SMARTseq, whereas 1,641 genes have significantly higher coverage in SoLo (paired T-test, p-adjusted < 0.05). Plotting the ratio of 5’ coverage between SoLo and SMARTseq similarly shows that as gene length increases, SoLo tends to have better coverage of the 5’ end than SMARTseq (Linear model: 0.872* log10(Length) - 2.084, R^2^ = 0.073, p-value < 0.001) (Figure 4E).

To determine whether these differences in length and coverage translate to differences in GeTMM levels, we compared log fold change values between SoLo and SMARTseq of the shortest 2000 mRNAs to the longest 2000 mRNAs, and found that the shortest genes had a wide distribution centered close to zero (median log2 fold change = −0.118), and the longest genes had a narrower distribution with most genes showing higher values in SoLo (median log2 fold change = 0.68) (p < 0.001) (Figure 4F). The relationship between dispersion and gene length is also different in SoLo and SMARTseq. Plotting gene level dispersion against average intra-technique GeTMM values reveals that gene dispersion is much lower in SoLo samples in the longest mRNAs (Supplementary Figure 3). The longest 2000 genes are overrepresented in genes called “expressed” in SoLo but “not expressed” in SMARTseq (2.5x expected), and underrepresented in the reverse comparison (0.27x expected). These results suggest that the longest genes make up ~20% of the SoLo exclusive genes. Together these data demonstrate that SoLo library preparation method results in both higher expression and better detection for longer genes.

### SoLo and SMARTseq show strong overlap with previous pan-neuronal gene sets

We benchmarked the two techniques against a published *C. elegans* pan-neuronal dataset to assess how well they replicate previous results. The Kaletsky dataset includes 8,437 protein coding genes called expressed in *C. elegans* neurons (3). Although the Kaletsky dataset was also derived from FACS-enriched neurons, the starting strain and the library construction methods differ (see Supplementary Table 2). Of the 8,437 Kaletsky expressed genes, 5,215 (61.8%) were called “expressed” in SMARTseq (Figure 5A). Of the remaining 3,222 genes called expressed in Kaletsky, 617 were called “not expressed” in SMARTseq, while another 2,605 (30.9%) were ambiguous. SoLo called 6,231 (73.9%) of the Kaletsky expressed genes “expressed” (Figure 5B). Of the remaining 2206 Kaletsky expressed genes, 456 genes were called “not expressed” in SoLo, while 1,740 genes (20.6%) were ambiguous. Thus, both techniques have broad agreement with previous data, and also show significant differences.

**Figure 5:**
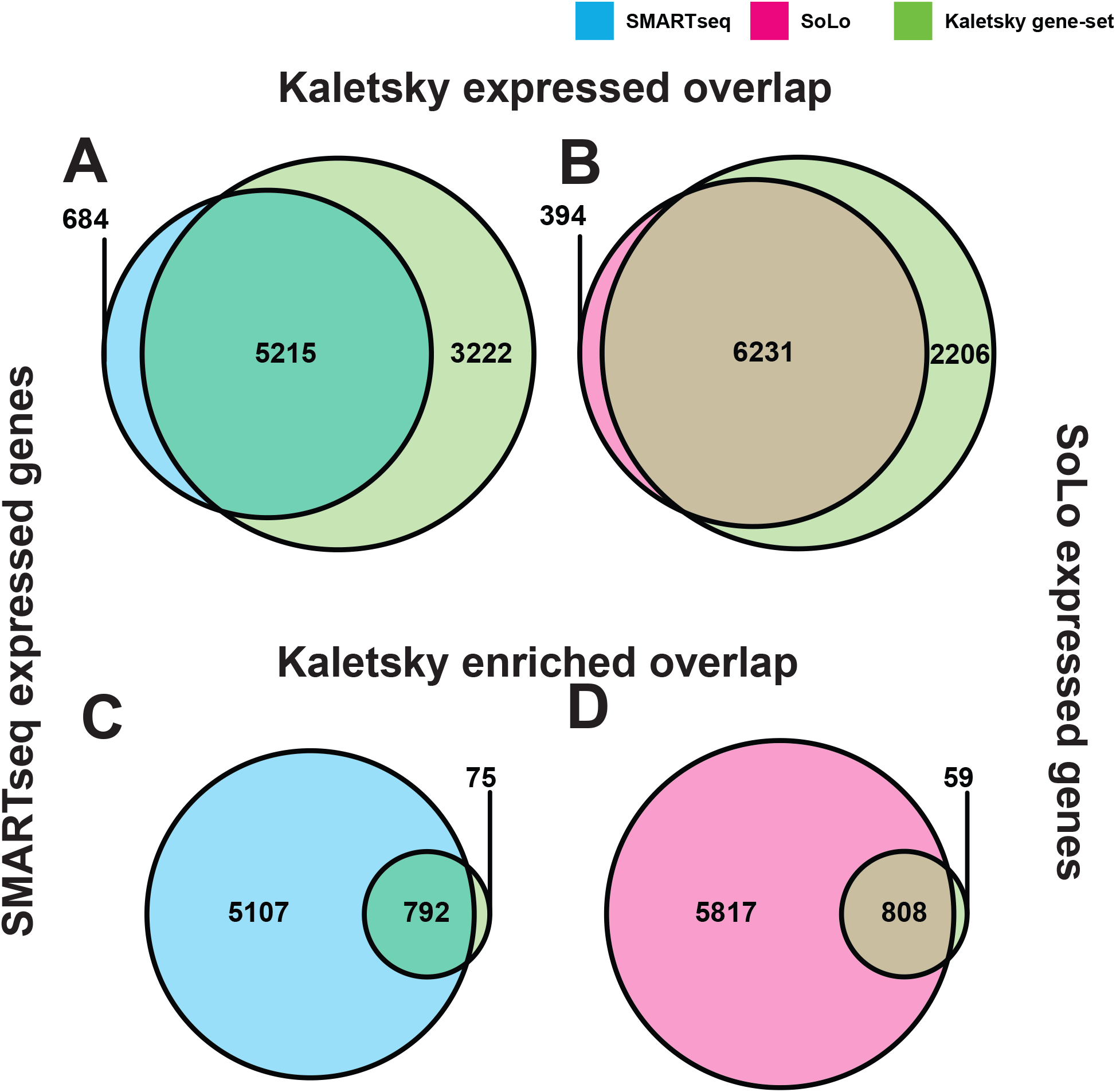
SoLo and SMARTseq detect neuronal genes. A-B) Overlap of “expressed” protein coding genes (lower bound of CI > 5 GeTMM) between SMARTseq (blue) or SoLo (magenta) and Kaletsky gene-set neuronal expressed genes (green). C-D) Overlap of “expressed” protein coding genes (lower bound of CI > 5 GeTMM) between SMARTseq (blue) or SoLo (magenta) and Kaletsky gene-set neuronal enriched genes (green).

The Kaletsky dataset also defines 867 neuronal enriched genes; these genes were not only found to be expressed in neurons but found to be enriched in neurons when compared to muscle, hypodermis, and intestinal cells. Of these, 792 (91.1%) were called “expressed” in SMARTseq (Figure 5C), and 808 (92.9%) were called “expressed” in SoLo (Figure 5D). These data show that >90% of neuronal protein coding genes in the Kaletsky gene set are called either “expressed” or “ambiguous” in both SoLo and SMARTseq, with minor differences explained by a mix of slight differences in contamination from other cell types during FACS, and different experimental parameters (Supplementary Table 2). Thus, our data from SoLo and SMARTseq approaches appears to strongly replicate previous findings for neuronal gene enrichment.

## DISCUSSION

Here we performed a head to head comparison of ribodepletion and polyA selection approaches for RNAseq library preparation using low input samples from *C. elegans*. Using RNA from FACS-isolated neurons, we evaluated the performance of SoLo Ovation (Tecan Genomics) and SMARTseq V4 (Takara) library preparation methods for rRNA depletion efficiency, overall library complexity and gene detection. Our results indicate that both techniques efficiently removed rRNA from the final libraries, although SMARTseq performed better than SoLo. SoLo libraries had fewer PCR duplicates than SMARTseq and detected more reads in UTRs. It is somewhat surprising that SoLo libraries contained fewer putative PCR duplicates than SMARTseq libraries from the same RNA inputs, considering that SoLo libraries also produced an order of magnitude more cDNA (Supplementary Table 1). This effect may be due to differences in where rRNA depletion occurs in the protocol. In the SMARTseq protocol, rRNA is selected against in the first step through preferential cDNA synthesis from polyA RNA. Downstream steps amplify only the targeted molecules across 16-20 rounds of PCR depending on the sample input, and any stochastic over-amplification will directly affect the targeted RNA species. For SoLo samples, all RNA is reverse transcribed to cDNA prior to initial amplification. In these initial rounds of PCR amplification, as rRNA sequences comprise the bulk of the cDNA, they are more likely to be represented in over-amplified products. Selected cleavage of rRNA adapters targets sequences that are already prone to stochastic over-amplification by virtue of their abundance. Thus, rRNA depletion after cDNA amplification in SoLo may partially protect against duplicate reads dominating the library during amplification of low input samples.

Noncoding RNAs play critical roles in gene regulation and cell function and detecting ncRNAs is key to fully understanding the transcriptome of any cell type. As many noncoding RNAs lack polyA tails, they may differ substantially in detection between SMARTseq and SoLo techniques. As expected, poly-adenylated noncoding RNAs, lincRNAs and pseudogenes, are detected at least as well by SMARTseq as they are by SoLo. Four classes of noncoding RNAs lacking polyA tails are detected at much higher rates in SoLo than SMARTseq. SoLo detects 13 times more tRNAs than SMARTseq, nearly seven times as many snRNAs, and 14 times as many snoRNAs, and 1.5 times as many uncategorized ncRNAs than SMARTseq, with a confidence interval above 5 GeTMM. tRNAs present their own challenges in sequencing, given their highly modified structure that often impedes RT-PCR. The robust detection seen here and the stark difference shown between approaches suggests that SoLo may be better suited to tRNA detection than SMARTseq, although further experiments are needed to confirm this finding. Overall, our results show that SoLo outperforms SMARTseq at detecting noncoding RNA species.

Given that each approach should theoretically treat poly-adenylated protein coding transcripts roughly the same, we set out to investigate whether these RNAs were detected at the same rate. The majority of protein coding genes called “expressed” in SoLo samples were also called “expressed” in SMARTseq, however SMARTseq was more prone to calling genes “ambiguous” than SoLo. We explain this difference by observing that estimated gene level dispersion is markedly higher among lowly expressed genes in SMARTseq compared to SoLo. This result suggests that SoLo may provide more confidence in calling genes expressed or not expressed, especially for genes that are expressed at low values.

While noise appears to drive much of the difference in the confidence of gene expression calling, we also investigated whether gene length bias drove differences in protein coding gene detection between the techniques. In studying the length distribution for high confidence and exclusive genes for each technique, we found that SMARTseq shows no clear deviation from the distribution of all protein coding transcripts, but SoLo shows a ~700bp increase in median transcript length. This finding corresponds with data on the average gene body coverage which shows SoLo having much more uniform coverage, especially at the 5’ end of the gene. On the basis of these findings, we hypothesized that long transcripts may be especially prone to reduced 5’ coverage in the polyA SMARTseq approach. The longer the gene, the more opportunity for degradation that severs the 5’ end from the 3’ polyA priming site. Additionally, longer genes may be more vulnerable to losing coverage of the 5’ end if the reverse transcriptase enzyme falls off of the RNA molecule prior to reaching the end. To test this idea, we measured coverage of the 5’ section of each gene and found that, for genes detected in both techniques, the ratio of SoLo coverage to SMARTseq coverage tends to increase with transcript length. By comparing the edgeR log fold changes for the shortest and longest protein coding transcripts we also found that while the shortest genes showed a wide range of fold changes centered close to zero, the longest genes were primarily enriched in SoLo, and were overrepresented in genes detected in SoLo and not detected in SMARTseq. Taken together these results suggest that expression of longer genes is generally prone to being underestimated, and that ribodepletion based techniques like SoLo are less vulnerable to this deficit. This disparity is unlikely to affect comparisons that focus solely on relative expression of a given transcript between conditions. However, the relative abundance of longer vs shorter genes within each condition could be underestimated due to this bias.

Kaletsky, et al. in 2018 published protein coding gene expression and enrichment lists for several *C. elegans* tissues, including a pan-neuronal dataset using an *unc-119* fluorescent reporter. This data set provided an opportunity to assess how well our SoLo and SMARTseq *rab-3* pan-neuronal libraries reproduce previous results. The comparison showed substantial overlap of SoLo/SMARTseq “expressed” protein coding genes with the Kaletsky dataset (Figure 5A-B), similar to our comparisons between the two library preparation techniques which use identical RNA samples (Figure 3A). Other differences between the Kaletsky data set and our results could be due to different fluorescent markers; the use of animals at different developmental stages; and differences in library preparation and gene thresholding procedures.

Overall, our findings suggest that SoLo Ovation, using a custom probe set to deplete *C. elegans* rRNA, outperforms SMARTseq with ultra-low input RNAseq samples by detecting an expanded set of noncoding RNAs, providing reduced noise for lowly expressed genes, and more accurate counts for long genes. Application of this technique, for example in efforts to profile all *C. elegans* neurons (27), should result in increased knowledge of cellular expression of diverse RNA molecules.

## METHODS

### Sample collection

The wild-type N2 strain and OH10689 *otIs355 [rab-3(prom1)::2xNLS-TagRFP]* IV were used for this study. Worms were grown on 8P nutrient agar 150 mm plates seeded with *E. coli* strain NA22. To obtain synchronized cultures of L4 worms, embryos obtained by hypochlorite treatment of adult hermaphrodites were allowed to hatch in M9 buffer overnight (16-23 hours) and then grown on NA22-seeded plates for 45-48 hours. The developmental age of each culture was determined by scoring vulval morphology (>75 worms) (28). Single cell suspensions were obtained as described (1, 3, 29) with some modifications. Worms were collected and separated from bacteria by washing twice with ice-cold M9 and centrifuging at 150 rcf for 2.5 minutes. Worms were transferred to a 1.6 mL centrifuge tube and pelleted at 16,000 rcf for 1 minute. 250 μL pellets of packed worms were treated with 500 μL of SDS-DTT solution (20 mM HEPES, 0.25% SDS, 200 mM DTT, 3% sucrose, pH 8.0) for 2 minutes.

Following SDS-DTT treatment, worms were washed five times by diluting with 1 mL egg buffer and pelleting at 16,000 rcf for 30 seconds. Worms were then incubated in pronase (15 mg/mL, Sigma-Aldrich P8811, diluted in egg buffer) for 23 minutes. During the pronase incubation, the solution was triturated by pipetting through a P1000 pipette tip for four sets of 80 repetitions. The status of dissociation was monitored under a fluorescence dissecting microscope at 5-minute intervals. The pronase digestion was stopped by adding 750 μL L-15 media supplemented with 10% fetal bovine serum (L-15-10), and cells were pelleted by centrifuging at 530 rcf for 5 minutes at 4 C. The pellet was resuspended in L-15-10, and single-cells were separated from whole worms and debris by centrifuging at 100 rcf for 2 minutes at 4 C. The supernatant was then passed through a 35-micron filter into the collection tube. The pellet was resuspended a second time in L-15-10, spun at 100 rcf for 2 minutes at 4 C, and the resulting supernatant was added to the collection tube.

Fluorescence Activated Cell Sorting (FACS) was performed on a BD FACSAria™ III equipped with a 70-micron diameter nozzle. DAPI was added to the sample (final concentration of 1 μg/mL) to label dead and dying cells. Cells were sorted under the “4-way Purity” mask. Sorted cells were collected directly into TRIzol LS. At ~15-minute intervals during the sort, the sort was paused, and the collection tube with TRIzol was inverted 3-4 times to ensure mixing. Cells in TRIzol LS were stored at −80° C for RNA extractions (see below).

### RNA Extraction

Cell suspensions in TRIzol LS (stored at −80° C) were thawed at room temperature. Chloroform extraction was performed using Phase Lock Gel-Heavy tubes (Quantabio) according to the manufacturer’s protocol. RNA in the aqueous layer was cleaned and concentrated using the RNA Clean and Concentrator Kit (Zymo Research, R1013). The aqueous layer from the chloroform extraction was combined with an equal volume of 100% ethanol and transferred to a Zymo-Spin IC column. Columns were centrifuged for 30 sec at 16,000 rcf. Samples 10 and 12 were then treated in-column with DNase I for 15 min (Supplementary Table 1). Samples 1 and 11 were not treated with DNase I. All samples were then washed with 400 μL of Zymo RNA Prep Buffer and centrifuged for 16,000 rcf for 30 sec. Columns were washed twice with Zymo RNA Wash Buffer (700 μL, centrifuged for 30 sec, followed by 400 μL, centrifuged for 2 minutes). RNA was eluted by adding 15 μL of DNase/RNase-Free water to the column filter and centrifuging for 30 sec. A 2 μL aliquot was submitted for analysis using the Agilent 2100 Bioanalyzer Pico chip to estimate yield and RNA integrity and the remainder stored at −80° C.

### rRNA probe optimization

To generate a probe set that targets *C. elegans* rRNA sequences, fasta sequences of all *C. elegans* rRNA genes were downloaded from wormmine (version WS235). Any exact duplicate sequences longer than 60 bases were reduced to a single copy. Tecan Genomics used these sequences to generate a set of 200 probes, proprietary to and available from, Tecan Genomics. Most rRNA genes were well covered, with the exception of a 150bp A/T rich region of MTCE.33 and a 400 BP A/T region at the 3’ end of MTCE.7.

### Library preparation and sequencing

SoLo and SMARTseq libraries were constructed for all samples according to manufacturer’s instructions. RNA concentrations are provided in Supplementary Table 1.

SoLo samples are treated with DNase I prior to first strand cDNA synthesis with a mix of oligo d(T) and random primers. First strand cDNA then undergoes hydrolysis, fragmentation, and strand selection before second strand synthesis, adapter ligation, Ampure bead purification, and amplification of the cDNA library with another round of Ampure bead purification. The amplified library is then incubated with the rRNA probe set, and rRNA fragments are selected against by nuclease-mediated cutting of the adapter sequence, followed by a final round of amplification and a final Ampure bead cleanup. Adapters in the Tecan SoLo Ovation kit use a single 8 base index, followed by an 8 base unique molecular identifier sequence (UMI).

SMARTseq samples are amplified using oligo d(T) primers to create the first strand of cDNA. A primer binding sequence is then appended to the first strand to allow for template switching and second strand synthesis. Long Distance PCR (LD-PCR) is then used to amplify the full-length cDNA into the final library before Ampure bead cleanup. cDNA libraries were then processed using the Illumina Nextera XT kit to create the sequencing library. Samples first underwent fragmentation and adapter ligation by Tn5 enzymatic tagmentation before final amplification and Ampure bead cleanup. Nextera uses dual-barcode adapters, but lack a UMI sequence.

All samples were checked for adapters dimers on an Agilent Bioanalyzer using the High Sensitivity DNA chip. Libraries were sequenced to a depth of 15-32 million read pairs on an Illumina HiSeq2500 machine, with paired end 75bp reads. SoLo libraries sequences on HiSeq4000 machines experienced run failures when not mixed with ~40% non-SoLo cDNA (data not shown). HiSeq2500 runs worked optimally when multiplexed samples were run at 5% higher concentration than standard.

### Read mapping, deduplication, sub feature base counting, and gene body coverage

Reads were mapped to the WBcel235 reference genome assembly using STAR (version 2.7.0). Duplicate reads were marked and removed using SAMtools (version 1.9.0). Counts files were generated using the featureCounts program from the SubRead package (version 1.6.4). Genes encloding rRNAs were removed from counts files prior to downstream normalization and gene detection steps.

Bases that map to exons, UTRs, introns, and intergenic regions were calculated using the CollectRNAseqMetrics program in PICARD (version 2.23.8).

Gene body coverage curves were obtained using the geneBody_coverage tool in RSeQC (version 2.6.4). To compare the coverage of the 5’ end between techniques, only protein coding genes called expressed in both techniques were considered. Gene models were split and each gene’s coverage was run separately. Within each gene, coverage was normalized to the highest coverage value. Average coverage across the first 20% of each gene was calculated for all replicates, and then averaged within each library building technique. Significance in 5’ coverage was calculated using a paired t-test in Scipy (version 1.5.0), p-values were adjusted for multiple hypothesis testing using the Benjamini-Hochberg correction.

### Gene expression normalization, thresholding, dispersion, and transcript length analysis

Counts matrices were normalized first to transcript length, and then to library size adjusted to the Trimmed-Mean of M (TMM) using edgeR (version 3.28.1) to obtain GeTMM values. 95% Confidence Intervals (CI) were calculated within each technique using the CI function in gmodels package in R (version 2.18.2). Normalized CI size was calculated for each gene by dividing the size of the interval by the Average GeTMM value (separately within each technique) to allow for easier comparison of CI sizes between genes that are lowly and highly expressed. Genes were called “not expressed” if the CI upper bound was < 5 GeTMM, genes were called “ambiguous” if the CI overlapped 5 GeTMM, and genes were called expressed if the CI lower bound was > 5 GeTMM. Dispersion estimates were generated for each library approach in edgeR using only samples from that library. Genes < 0.5 average GeTMM showed identical gene level dispersion and were excluded. Differential expression analysis was calculated using a generalized linear model in edgeR. Analyses that focused on protein coding gene expression used only protein coding genes to calculate TMM values and library size. Transcript lengths were used from the output of featureCounts. Pseudocounts of 1 were added for all log10 GeTMM plots. When calculating ratios of 5’ coverage, gene dispersion, or normalized CI size, pseudocounts of 1*10^^−5^ were added to the numerator and denominator to avoid infinite values without unduly skewing results visually.

## Data Availability

Raw fast files and a processed counts table have been submitted to GEO, and this manuscript will be updated with the appropriate accession number when available. In the meantime, authors will provide fastq and counts files directly upon request.

---

**Supplementary figure 1:**
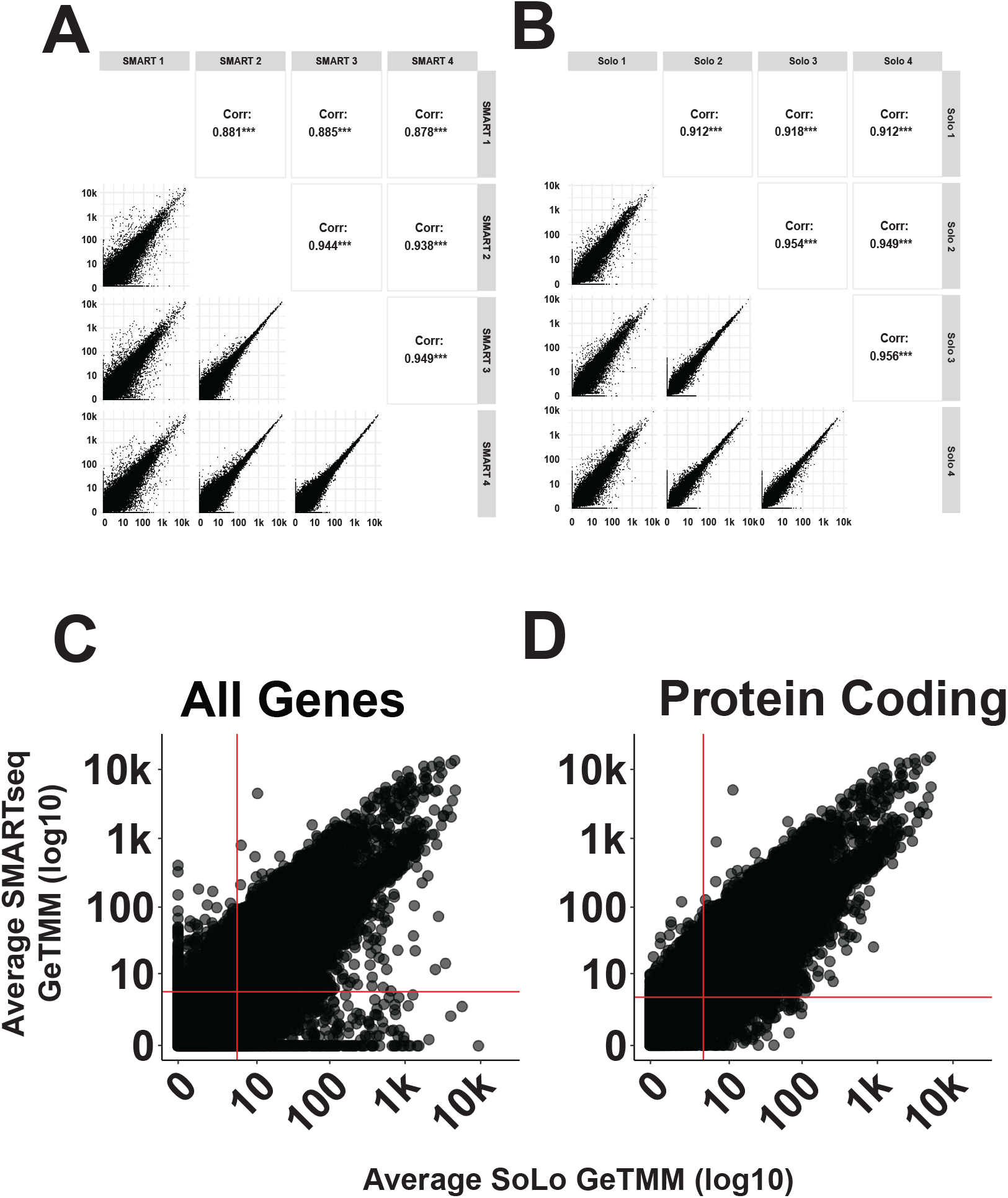
GeTMM values correlate within techniques. A-B) Scatter plots with correlation values for all pairwise replicates within SMARTseq (A) and SoLo (B) samples. C-D) Scatter plots showing averaged log10 GeTMM values for SoLo and SMARTseq samples using all genes (C), and just protein coding genes (D).

**Supplementary figure 2:**
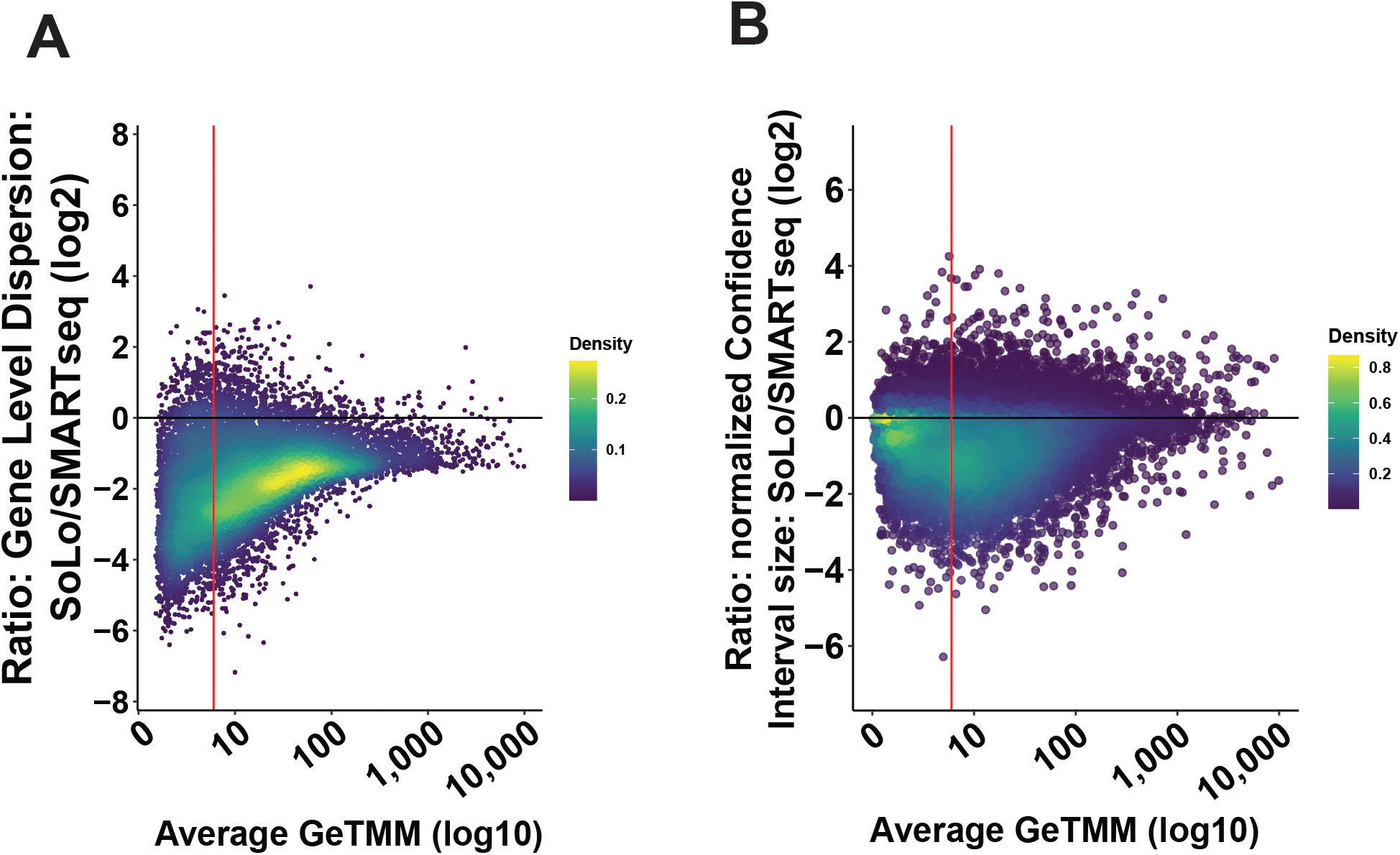
SMARTseq samples show higher noise on gene level. A) Scatterplot showing log2 ratio of edgeR gene level dispersion estimate between SoLo and SMARTseq against average GeTMM value. Data points are colored by density on the graph. B) Scatterplot showing log2 ratio of expression normalized confidence interval size between SoLo and SMARTseq against average GeTMM value. Data points are colored by graph density.

**Supplementary figure 3:**
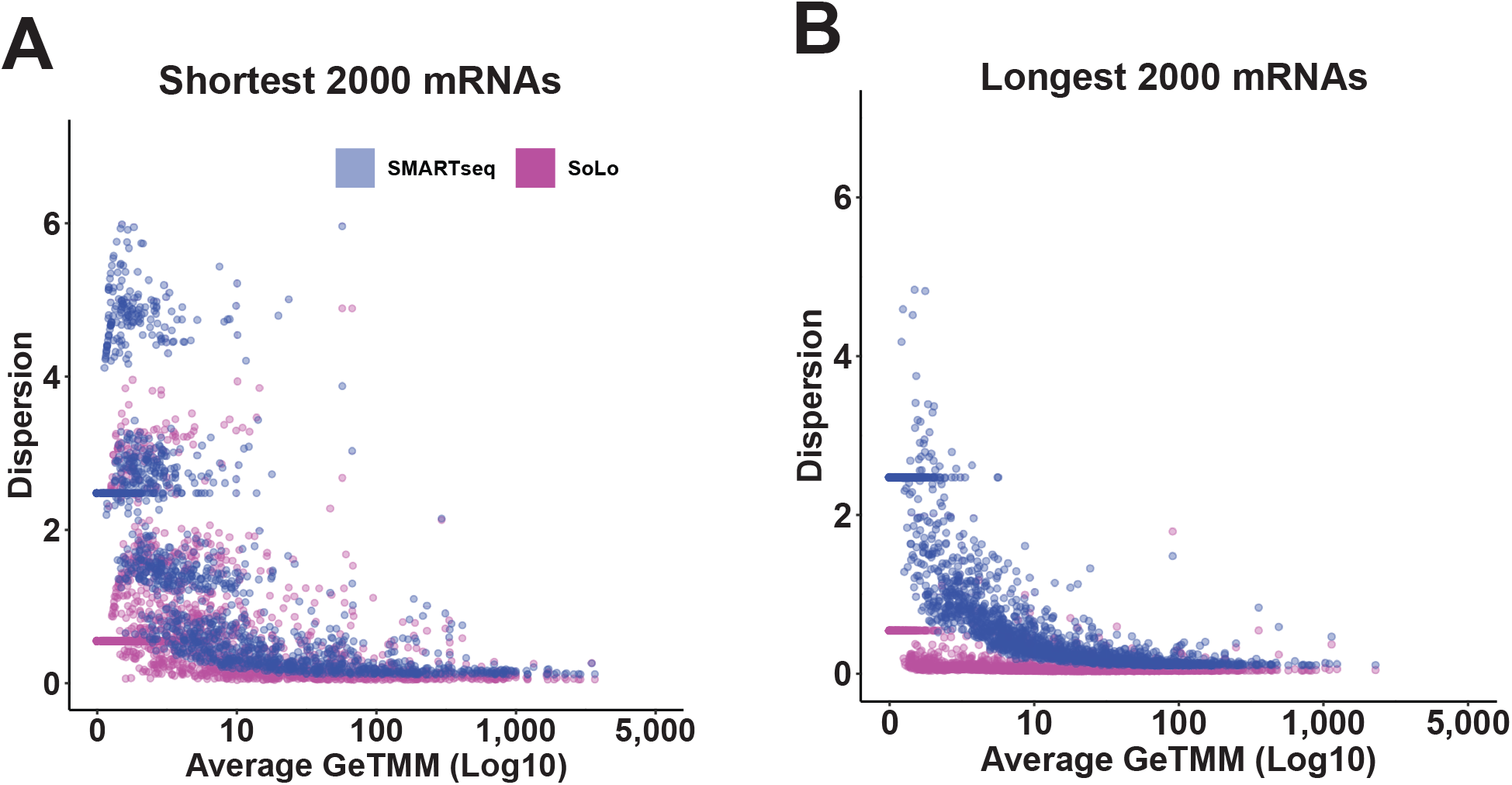
Difference in Dispersion between SoLo and SMARTseq is higher for longest genes. A-B) Scatter plot showing the edgeR gene level dispersion estimate against the average GeTMM levels for SMARTseq (blue) and SoLo (magenta). Plots show the shortest 2000 protein coding genes (minimum length > 100) (A), or the longest 2000 protein coding genes (B).

**Supplementary Table 1:** Sample metadata

| sample name | total rna (ng) | RIN | Total read pairs | Total Mapped | Total Mapped, deduplicated | % uniquely mapped | % rRNA reads | % rRNA after deduplication | DNase treated? | Biological replicate | Final library cDNA yields (ng) |
| --- | --- | --- | --- | --- | --- | --- | --- | --- | --- | --- | --- |
| SMARTseq 1 | 1.508 | 9.8 | 31767947 | 25142260 | 7935521 | 79.14348 | 2.505802 | 1.708252 | No | 1 | 3.68 |
| SMARTseq 2 | 1.160 | 9.5 | 40416985 | 25631426 | 5915352 | 63.41746 | 3.373189 | 2.315434 | Yes | 2 | 3.00 |
| SMARTseq 3 | 3.600 | 9.9 | 37442564 | 24398942 | 7394695 | 65.16365 | 3.553103 | 2.194880 | No | 3 | 4.36 |
| SMARTseq 4 | 1.230 | 9.8 | 36977605 | 23990669 | 5018616 | 64.87892 | 2.927278 | 2.223749 | Yes | 3 | 3.68 |
| SoLo 1 | 1.508 | 9.8 | 17969902 | 10483341 | 3860066 | 58.33833 | 22.203690 | 11.712460 | No | 1 | 59.20 |
| SoLo 2 | 1.160 | 9.5 | 17395388 | 10075674 | 3001022 | 57.92152 | 24.961320 | 15.558520 | Yes | 2 | 60.00 |
| SoLo 3 | 3.600 | 9.9 | 21456169 | 12216428 | 4393602 | 56.93667 | 25.743630 | 12.983270 | No | 3 | 78.00 |
| SoLo 4 | 1.230 | 9.8 | 15559476 | 9323351 | 3010153 | 59.92073 | 17.912890 | 12.886540 | Yes | 3 | 60.60 |

**Supplementary Table 2:** Comparison to Kaletsky et al., 2018 samples

| Dataset | Developmental stage | Fluorescent reporter used | RNA input range | cDNA synthesis kit | Sequencing library prep kit | Sequencing machine |
| --- | --- | --- | --- | --- | --- | --- |
| SoLo | L4 | otIs355<br>[rab-3(prom1)::2xNLS-TagRFP] | 1.5-3.6 ng | Tecan/Nugen SoLo ovation | Tecan/Nugen SoLo ovation | HiSeq 2500 |
| SMARTseq | L4 | otIs355<br>[rab-3(prom1)::2xNLS-TagRFP] | 1.5-3.6 ng | Takara SMARTseq V4 | Illumina Nextera XT | HiSeq 2500 |
| Kaletsky et al., 2018 (neuronal) | Day 1 Adult | otIs45<br>[Punc-119::GFP] | 10-100 ng | Tecan/Nugen Ovation RNAseq v2 | Tecan/Nugen Encore NGS OR Illumina TruSeq DNA Sample Prep | HiSeq 2000 |
| Kaletsky et al., 2018 (neuronal) | Day 1 Adult | otIs45<br>[Punc-119::GFP] | 10-100 ng | Takara SMARTer Stranded Total RNA kit-pico input mammalian | Takara SMARTer Stranded Total RNA kit-pico input mammalian | HiSeq 2000 |

## Acknowledgements

This work was funded by NIH grant R01NS100547 to MH, OH, DMM, and NS and by Vanderbilt Trans-Institutional Program funds to DMM. Flow Cytometry experiments were performed in the Vanderbilt Flow Cytometry Shared Resource which is supported by the Vanderbilt Ingram Cancer Center (P30 CA68485) and the Vanderbilt Digestive Disease Research Center (DK058404). The Vanderbilt VANTAGE Core provided technical assistance for this work and is supported by CTSA Grant (5UL1 RR024975-03), the Vanderbilt Ingram Cancer Center (P30 CA68485), the Vanderbilt Vision Center (P30 EY08126), and NIH/NCRR (G20 RR030956). Some strains were provided by the CGC, which is funded by NIH Office of Research Infrastructure Programs (P40 OD010440).

